# Massive culture-based approach for the screening of AmpC-, ESBL- and carbapenemase producers from complex samples

**DOI:** 10.1101/2025.01.06.631451

**Authors:** Gabriel Taddeucci-Rocha, Victoria de Oliveira Costa, Sarah Vitória Martins da Silva, Jéssica Britto Gonçalves, Natalia Chilinque Zambão da Silva, Marcia Maiolino Garnica, Renata Cristina Picão

## Abstract

Gram-negative bacilli producing beta-lactamases are major causes of difficult-to-treat infections, especially the AmpC, ESBL, and carbapenemase types. Their spread within and outside hospital settings demands effective detection and monitoring in various environments, but current methods for this purpose often neglect important groups of beta-lactamases or are expensive and time consuming. We aimed to develop and test a massive culture approach to detect and differentiate between beta-lactamase producers from complex samples. The method involved an enrichment step on MacConkey agar supplemented with ceftriaxone to select for AmpC, ESBL, and carbapenemase producers, followed by replica plating under selective pressures (cefoxitin, cefepime, imipenem) to differentiate them. The massive culture approach effectively differentiated strains producing different beta-lactamases in mixed cultures. In tests with rectal swabs, our method demonstrated 100% sensitivity, higher specificity, and greater accuracy for ESBL detection compared to the reference method. Additionally, it identified a broader spectrum of beta-lactamase producers, including AmpC and carbapenemase. The massive culture approach is a promising tool for detecting and differentiating gram-negative bacilli producing beta-lactamases from rectal swabs. Due to the additional time required to produce results, this method is most suitable for central and research laboratories and enhances surveillance capabilities for antimicrobial resistance.

**IMPORTANCE:** The dissemination of multidrug-resistant bacteria producing beta-lactamases in healthcare and community settings demands effective detection and monitoring. Existing methods may miss key beta-lactamase groups or are costly and time-consuming. In this study, a comprehensive culture-based approach using replica plating was developed and used to detect and differentiate beta-lactamase producers from rectal swabs. It showed higher specificity and greater accuracy for ESBL detection compared to the reference method, while identifying a broader range of beta-lactamase producers, including AmpC and carbapenemase. Despite requiring additional processing time, this method is suitable for central and research laboratories, enhancing antimicrobial resistance surveillance.

## INTRODUCTION

Beta-lactam-resistant gram-negative bacilli are major causes of infections in both hospitals and community settings, primarily due to the production of beta-lactamases (1). These enzymes include AmpC-like cephalosporinases, extended-spectrum beta-lactamases (ESBL), and carbapenemases, which confer resistance to various beta-lactams (2). Genes encoding these beta-lactamases are often carried along with resistance determinants against other antimicrobial classes, severely limiting therapeutic options (3). For these reasons, beta-lactamase producers are key targets for antimicrobial resistance (AMR) research (4).

The spread of beta-lactamase producers outside hospitals is a growing global concern with unknown consequences for human, animal and environmental health (5). Thus, monitoring their presence in hospitals is crucial for infection control, while surveillance of carriage in asymptomatic humans, animals, and environments is essential for risk assessment and understanding the environmental spread of AMR (6).

Surveillance of these microorganisms among humans typically involves culturing fecal specimens under selective pressure to target specific mechanisms such as ESBL or carbapenemase production, followed by analysis of isolated colonies. Molecular detection methods for beta-lactamase resistance mechanisms exist but often focus on a few known carbapenemases, neglecting acquired ESBLs, AmpCs, some carbapenemases and undescribed beta-lactamase families that may also threaten anti-infective therapy (7); or involves sequencing of the entire resistome using metagenomics (8). The latter requires specialized equipment and highly trained personnel in bioinformatics, limiting its use for routine diagnostic and research purposes, especially in low- and middle-income countries (9). Consequently, phenotypic investigation through selective culture remains the reference method to survey carriage of beta-lactamase producers. However, colonization may involve different strains carrying various AMR mechanisms, so the targeted culture-based approach may limit the understanding of strains and mechanisms driving the spread of beta-lactam resistance in different settings (10).

Replica plating is a classical method that allows the evaluation of all colonies present in a culture and enables assessment of their growth under various selective pressures (11). We hypothesized that a massive culture approach based on replica plating is suitable for detecting and differentiating beta-lactamase producers from complex samples. Our study aims to: i) standardize a massive culture approach for the identification of AmpC, ESBL and carbapenemase producers; ii) evaluate its performance in detecting ESBL producers in rectal swabs; and iii) describe the detection of plasmid-mediated AmpC (pAmpC) and/or carbapenemases in rectal swabs using this approach.

## MATERIALS AND METHODS

### Protocol standardization

The approach we envisioned is based on a enrichment culture on MacConkey agar supplemented with ceftriaxone, followed by replica plating under selective pressures using cefoxitin, cefepime and imipenem to differentiate between AmpC, ESBL and carbapenemase producers, respectively (Supplementary material, Fig. S1).

Initially, we aimed to identify the ideal drug concentration for replica plating by testing three gram-negative bacilli that produce different beta-lactamases (pAmpC, ESBL and carbapenemase) and exhibit distinct colony characteristics on MacConkey agar, along with a negative control. These included ESBL-producing *Klebsiella pneumoniae* 134 (CTX-M-15, large mucoid lactose-fermenting colonies) (12), plasmid-mediated AmpC-producing *Escherichia coli* D493 trans-III (CMY-2, medium-sized opaque lactose-fermenting colonies) (13), and carbapenemase-producing *Pseudomonas aeruginosa* C237 (VIM-2, medium-sized flat colonies with irregular margins, lactose-non-fermenting) (14), and the negative control *E. coli* TOP10 (small lactose-non-fermenting colonies).

An initial inoculum of 10^8^ CFU/ml was prepared for each bacterium using the 0.5 McFarland standard as reference. Equal volumes of each suspension were combined in a single 1,5-ml tube and diluted to reach 10^6^ CFU/mL and 10^5^ CFU/mL. Fifty-microliter aliquots of the diluted mixed suspensions were seeded on antimicrobial-free MacConkey agar plates without antimicrobials (MAC), and on MAC supplemented with ceftriaxone (MCRO, 1.5 µg/ml), in triplicate, and incubated overnight at 35 ± 2 °C under aerobic conditions. The microbial growth on each plate was replicated onto eight MAC: one initial and one final plate without antimicrobials, for inoculum control; and two plates with each drug concentration (1x and 2.5x the breakpoint for resistance according to the CLSI M100-Ed33 document). These included cefepime (FEP, Sigma-Aldrich, 16 and 40 µg/ml), cefoxitin (FOX, Sigma-Aldrich, 32 and 80 µg/ml), and imipenem (IPM, Sigma-Aldrich, 4 and 10 µg/ml).

Sterile 15x15 cm pieces of 2.0mm-thick elastic velvet were fixed to an 89 mm-diameter support made of autoclavable resin and printed on an SLA 3D printer. Velvet sterile control was assessed by pressing a tryptic soy agar (TSA) plate against the fabric. Fresh cultures from MCRO were transferred to the velvet by gently pressing the culture against it, followed by replication on the series of plates by gently pressing each plate against the velvet. All culture plates were incubated overnight at 35 ± 2 °C under aerobiosis. The parameter to choose the best concentration to be used in further experiments was observing well-formed colonies corresponding to the expected microorganism(s): in plates supplemented with cefepime it should only grow those strains producing ESBL and carbapenemase; in plates supplemented with cefoxitin it should only grow strains producing AmpC and carbapenemase; and in imipenem it was only expected the growth of strains producing carbapenemase (Fig. S1).

We then assessed the performance of the massive culture approach in a non-controlled sample. For this, we seeded a rectal swab provided by a volunteer in MCRO and replicated the culture onto five MacConkey agar plates: without supplementation, with cefepime (MFEP, 16 µg/mL), with cefoxitin (MFOX, 32 µg/mL), with imipenem (MIPM, 4 µg/mL), and again without supplementation. The detailed protocol is available (supplemental material and supplemental video). Colonies observed on MFOX, MFEP and MIPM were investigated for plasmid-mediated Ampc (pAmpC), ESBL and carbapenemase production, respectively, as described below.

### Accuracy of the massive culture approach to detect ESBL producers in rectal swabs

We evaluated the performance of the proposed method for detecting ESBL in self-collected rectal swabs, comparing it to the performance of a commercial chromogenic agar (Laborclin, PR, Brazil), hereafter referred to as CESBL. The fecal specimens were provided by 30 volunteers. The swabs were first spotted in TSA to verify the presence of microorganisms, ensuring that the specimens were properly collected. They were then cultured on CESBL and MCRO and incubated aerobically during 18–24 hours at 35 ± 2 °C. Specimens showing no bacterial growth on TSA were excluded from further analysis. To avoid biased results due to insufficient fecal material, we first cultured 15 swabs on CESBL and then on MCRO; while another 15 swabs were first inoculated on MCRO and then on CESBL.

MCRO plates showing bacterial growth were replicated onto five MacConkey agar plates: MAC, MFOX, MFEP, MIPM, MAC. For the specific purpose of evaluating the accuracy of the massive culture approach in detecting ESBL producers, only colonies grown on CESBL and MFEP were further studied. Up to five representatives of each colony morphotype were selected and identified using MALDI-TOF (Bruker). Enterobacteria were evaluated for ESBL production by the double disk synergy test, and positive isolates were confirmed by PCR targeting *bla*CTX-M-1/2-like, *bla*CTX-M-8-like, *bla*CTX-M-9-like, *bla*SHV-like, *bla*TEM-like and *bla*GES-like genes. *Pseudomonas* spp. and *Acinetobacter* spp. were only investigated for the presence of ESBL-encoding genes due to the poor performance of the phenotypic test in detecting ESBL production in these microorganisms. Isolates where ESBL production is clinically irrelevant were not investigated for this resistance mechanism.

The growth of colonies with confirmed ESBL production was considered a true-positive result for both CESBL and the massive culture approach on MFEP. False-positives were defined as the growth of colonies without such confirmation. True-negative results were recorded when no colonies grew on selective media for both tests, and false negatives were noted when there was no growth in one technique but a true-positive result in the other. Sensitivity, specificity, positive predictive values (PPV), and negative predictive values (NPV) were calculated to compare the performance of the massive culture approach and the reference method in detecting ESBL producers in rectal swabs (Table S1, supplement material).

### Detection of pAmpC and carbapenemase producers using the massive culture approach

We additionally studied the colonies grown under cefoxitin and imipenem selective pressures from rectal swabs to assess the occurrence of AmpC and carbapenemase producers, respectively. Up to five representatives of each colony morphotype were subcultured under the same conditions and identified using MALDI-TOF. Production of pAmpC was assessed among *E.coli* and *Klebsiella* spp., known as susceptible to third generation cephalosporins due to low expression or absence of intrinsic AmpC-like cephalosporinases, respectively. Isolates resistant to cefoxitin and amoxicillin/clavulanate, assessed by CLSI disk diffusion (15), were subjected to PCR targeting genes encoding MOX, CIT, DHA, ACC, EBC and FOX pAmpC groups, as previously described (16), followed by amplicon sequencing. Carbapenemase production was evaluated using the modified carbapenem inactivation method (mCIM) (17, 18) and the EDTA-modified carbapenem inactivation method (eCIM) (19), and confirmed by the detection of carbapenemase-encoding genes (20).

### Ethics in research

This study was approved by the Human Research Ethics Committee (protocol number 46762621.2.0000.5455), and each participant signed an informed consent form.

## RESULTS

We first optimized the protocol by establishing the drug concentrations that would result in growth of well-formed colonies of the expected microorganisms, using previously characterized bacteria. We observed that the lowest tested concentration of all antimicrobials allowed only the expected bacteria to grow, while increased concentrations showed a partial inhibitory effect on the target bacteria, particularly in the case of cefoxitin. The growth observed in the last inoculum control plate resembled that of the initial one, ensuring that the growth inhibition in the intermediate plates was due to antimicrobial action and not to decrease in the bacterial load throughout the series of replicates (Fig. 1). It should be noted that we were able to detect the resistant strain on its corresponding selective pressure even when its inoculum was smaller and completely covered by other colonies in the initial culture (Fig. 1). Therefore, we selected the lower concentrations of antimicrobials for use to survey for beta-lactamase producers.

**Figure 1.**
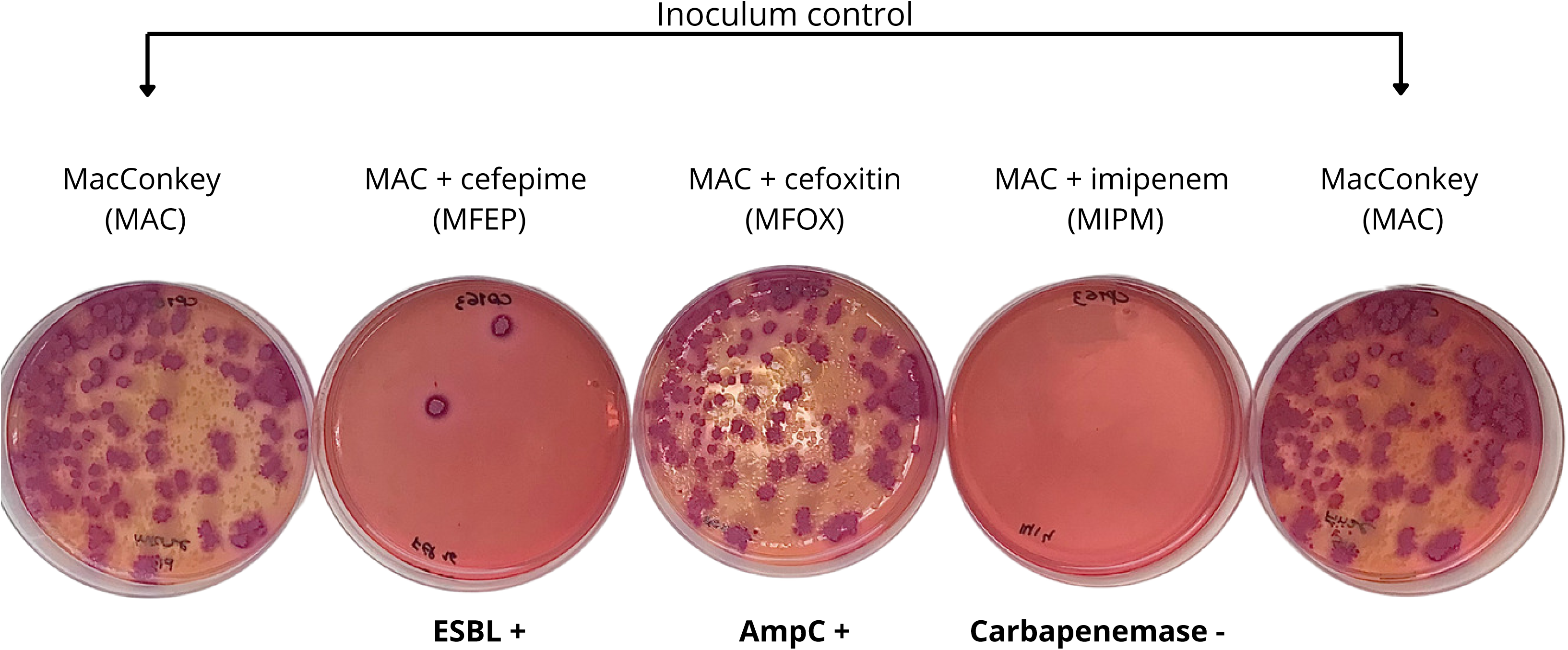
Result of massive culture using replica plating for different CFU/ml counts of a mixed suspension, including previously characterized strains producing pAmpC, ESBL, carbapenemase, and a negative control (*E. coli* TOP10). The suspensions were initially cultured overnight on MacConkey ágar without supplementation (MAC), and then replicated onto: MAC plate for initial inoculum control; cefoxitin (MFOX 32 and 80 µg/ml), cefepime (MFEP 16 and 40 µg/ml), imipenem (MIMP 4 and 10 µg/ml), and a MAC plate for final inoculum control. Growth of all strains is expected in MAC plates; while the CTX-M-15-producing *K. pneumoniae* (forming large, mucoid, and lactose-fermenting colonies) is expected to grow in MFEP; CMY-2-producing *E. coli* expected to grow on MFOX (forming medium-sized, opaque, lactose-fermenting colonies); and the VIM-2-producing *P. aeruginosa* is expected to grow on MFOX, MFEP and MIPM (forming medium-sized flat colonies with irregular margins, lactose-non-fermenting).

We subsequently tested the suitability of this approach using rectal swabs, with positive results for differentiating various beta-lactamases present, regardless of their differing counts in the clinical specimen. Figure 2 shows the resulting cultures from the replication of a MCRO plate inoculated with the specimen. Most colonies present in the control plates were also found in MFOX, suggesting production of AmpC. Notably, it was also possible to observe 3 likely ESBL-producing CFU in MFEP and complete growth inhibition on MIPM, suggesting absence of carbapenemase producers. Eight isolates obtained from MFOX were identified as *Citrobacter* spp. (4) and *Pseudomonas* spp. (4), thus intrinsic AmpC producers. The three colonies grown on MFEP were identified as *E. coli* (2). These isolates exhibited a positive ESBL phenotype that was confirmed by the amplification of the *bla*_CTX-M-79_ and *bla*_CTX-M-55_ genes for both *E. coli*, showing 100% identity for both genes, with an amplicon size of 839 bp.

**Figure 2.**
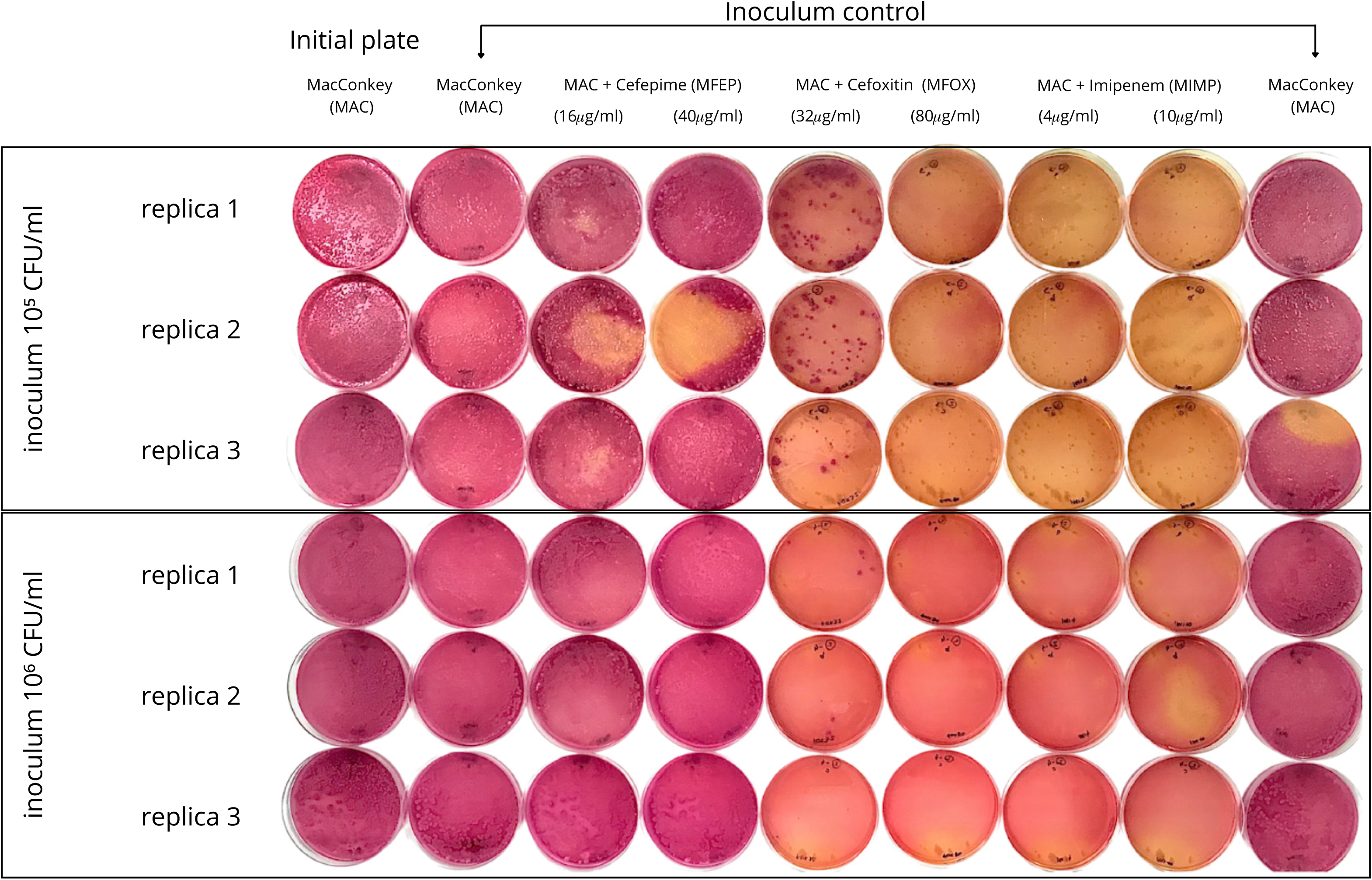
Replication of growth observed after seeding the rectal swab on MacConkey agar plate supplemented with ceftriaxone (1.5 µg/mL) on a series of MacConkey agar plates: without supplementation, with cefepime (MFEP, 16 µg/mL), with cefoxitin (MFOX, 32 µg/mL), with imipenem (MIPM, 4 µg/mL), and again without supplementation. Plates without antibiotics are intended for inoculum control, while plates with cefoxitin is intended to select for AmpC and carbapenemase producers, cefepime for ESBL and carbapenemase producers and imipenem for carbapenemase producers. The volunteer providing the swab showed colonization by an AmpC intrinsic producer (*Citrobacter* spp. and *Pseudomonas* spp.) and ESBL-producing *E. coli* confirmed by phenotypic and genotypic tests.

We compared the performance of the massive culture approach to detect ESBL producers with that of the reference method, using CESBL. We searched for gram-negative bacilli producing beta-lactamases in 25 rectal swabs (five specimens were excluded for showing no growth in the TSA control, thus were considered improperly collected) among which 10 were detected to carry ESBL producers by both techniques. Despite the agreement on the number of swabs with ESBL producers, the profiles observed, considering bacterial identification and ESBL-encoding genes, were not always concordant between CESBL and the massive culture replica under selective pressure by cefepime (Table S2). In total, 159 bacteria were isolated, with 71% identified as enterobacteria, 28% as non-fermenting gram-negative bacilli (BGN-NF), and 1% gram-positive cocci. Only the enterobacteria confirmed ESBL production. Cultures resulting from the replica plating in MFEP showed a higher percentage of Enterobacteriaceae (88%) and the highest percentage of ESBL producers among the total microorganisms recovered (67%), compared to the CESBL approach (60% enterobacteria and 37% of ESBL producers). The massive culture approach identified 81% of the ESBL producers profile, considering microbial identification and the ESBL-encoding genes amplified, whereas CESBL detected 74%. Moreover, the number of Enterobacteriaceae that were negative for ESBL production was higher in CESBL (22) than in MFEP (13) (Table S3). Similar findings were observed for *Acinetobacter* spp. and *Pseudomonas* spp., which showed negative results by PCR and were more frequently found in CESBL (17 and 18, respectively) than in MFEP (3 and 2, respectively) (Table S4). The sensitivity for both techniques was 100%, however, the specificity was higher for the massive culture approach (20% vs 47%, respectively), which translated into greater accuracy than CESBL (68% vs 52%), respectively (Table S1).

We investigated colonies that grew on the MFOX and MIPM replicas to describe occurrences of bacteria producing pAmpC and carbapenemases, respectively. A total of 11 swabs showed bacterial growth on the MFOX replica, resulting in 39 isolates. Twenty-six isolates obtained from MFOX were identified as *Pseudomonas* spp. (15), *Enterobacter* spp. (5), *Citrobacter* sp. (3) and *Acinetobacter* sp. (2), thus intrinsic AmpC producers, and *Enterococcus* sp. (1), a gram-positive cocci. Among the remaining isolates, ten *E. coli* and three *K. pneumoniae* recovered from three and one volunteers, respectively, were surveyed for genes encoding pAmpC. Four *E. coli* isolates from volunteer 18 confirmed the presence of *bla*CMY-like; four isolates from volunteer 30 confirmed the presence of *bla*NDM-like; and two *E. coli* isolates from volunteer 2 were negative for pAmpC, but confirmed the genes *bla*TEM-like, *bla*CTX-M-8 and *bla*CTX-M1/2-like genes. Of notice, volunteer 2 also showed *E. coli* positive for the same genes in the MFEP replica (Table S2). No pAmpC-encoding genes were observed for the three *Klebsiella* isolates recovered from volunteer 3, but all of them showed *bla*SHV, *bla*TEM and *bla*CTX-M1/2-like genes, a genotype that was also observed in *Klebsiella* sp. recovered from the same subject in the MFEP replica (Table S2).

Three swabs showed bacterial growth on the MIPM replicas, resulting in 12 isolates. Isolates obtained from MIPM were identified as *E. coli* (6), *Enterobacter* spp. (2), *Klebsiella* sp. (2) and *Stenotrophomonas* sp. (2). Only the six *E. coli* recovered from the swab of volunteer 30 confirmed acquired carbapenemase activity that was inhibited by EDTA, and it amplified the *bla*NDM gene.

## DISCUSSION

The massive culture approach based on replica plating under selective pressure, designed to detect AmpC, ESBL, and carbapenemase producers in complex samples, proved capable of separating strains producing specific beta-lactamases even when they are outnumbered by other bacterial colonies. Its use in rectal swabs showed greater specificity than the targeted approach for detecting ESBL producers, with the added advantage of simultaneously identifying isolates producing AmpC and carbapenemases.

The method was designed with a two-step process: the first step selects beta-lactamase-producing bacteria, and the second step separates them. The most commonly used approach for ESBL production, CESBL, corresponds to our first step. The subsequent replica plating achieves the separation of different beta-lactamases, thus eliminating many intrinsic AmpC producers that yielded false positives in the reference method. This likely explains why we achieved higher specificity.

Another notable characteristic of the proposed method is its enhanced ability to identify different profiles of ESBL producers, considering the identification and the array of beta-lactamases produced. Even with a small set of clinical specimens, we were able to identify a significant number of profiles of Enterobacteriaceae producing various beta-lactamases, not only ESBL, but also pAmpC and the NDM carbapenemase. This is particularly significant given that colonization is a highly dynamic process (10). Using this method, we have a better chance of accessing this dynamism with a single clinical specimen without the risk of depleting it across different culture media. In practice, this minimizes the potential for biased results concerning the global colonization status of the gastrointestinal tract by beta-lactamase-producing bacteria.

The main disadvantage of the massive culture approach is that it takes an additional day to provide results compared to the targeted approach, which could be a drawback for infection control applications. Nonetheless, in an ideal scenario, all culture-based methods are more suited for central and research laboratories rather than for guiding bedside decision-making, where methods that provide results within a few hours are preferred.

Despite this limitation, our findings indicate that the proposed method can be advantageous for investigating Enterobacteriaceae producing various impactful beta-lactamases. Our results may encourage researchers to adopt this approach for studying these important microorganisms not only in clinical specimens but also in environmental samples; and to standardize the best concentrations for investigating resistance determinants to other antimicrobial classes.

## ACKNOWLEDGMENTS

We thank the volunteers that provided the rectal swab samples that viabilized the present work. We also thank the sponsors Conselho Nacional de Desenvolvimento Científico e Tecnológico (CNPq), Fundação Carlos Chagas Filho de Amparo à Pesquisa do Estado do Rio de Janeiro (FAPERJ); Instituto Nacional de Pesquisa em Resistência Antimicrobiana; and Coordenação de Aperfeiçoamento de Pessoal de Nível Superior. The sponsors had no role in the design and conduct of the study; collection, management, analyses, and interpretation of the data; preparation, review, or approval of the manuscript; and decision to submit the manuscript for publication.

## DATA AVAILABILITY STATEMENT

Data available on request due to privacy/ethical restrictions.

## FUNDING

This work was supported by Conselho Nacional de Desenvolvimento Científico e Tecnológico (CNPq) [process numbers 311737/2019-6, 408725/2022-2]; by Fundação Carlos Chagas Filho de Amparo à Pesquisa do Estado do Rio de Janeiro (FAPERJ) [process numbers, E-26/200.795/2019, E-26/211.554/2019, E-26/201.191/2021, E-26/211.351/2021]; by INPRA - Instituto Nacional de Pesquisa em Resistência Antimicrobiana - Brazil [INCT/CNPq: 465718/2014-0]; by Coordenação de Aperfeiçoamento de Pessoal de Nível Superior - Brazil [CAPES, financing code 001].

## CONFLICTS OF INTEREST

The authors declare no conflict of interest.

## CONTRIBUTION

GTR: Investigation, Methodology, Data curation, Formal analysis, Supervision, Writing - original draft, Writing - review & editing; VOC: Investigation, Methodology, Data curation, Writing - review & editing; SVMS: Investigation, Methodology, Writing - review & editing; JBG: Investigation, Methodology, Writing - review & editing; NZCS: Investigation, Supervision, Writing - review & editing; MG: Supervision, Writing review & editing; RCP: Conceptualization, Data curation, Formal analysis, Supervision, Writing review & editing.

## REFERENCES

1. Murray CJL, Ikuta KS, Sharara F, Swetschinski L, Aguilar GR, Gray A, Han C, Bisignano C, Rao P, Wool E, Johnson SC, Browne AJ, Chipeta MG, Fell F, Hackett S, Haines-Woodhouse G, Hamadani BHK, Kumaran EAP, McManigal B, Achalapong S, Agarwal R, Akech S, Albertson S, Amuasi J, Andrews J, Aravkin A, Ashley E, Babin F-X, Bailey F, Baker S, Basnyat B, Bekker A, Bender R, Berkley JA, Bethou A, Bielicki J, Boonkasidecha S, Bukosia J, Carvalheiro C, Castañeda-Orjuela C, Chansamouth V, Chaurasia S, Chiurchiù S, Chowdhury F, Donatien RC, Cook AJ, Cooper B, Cressey TR, Criollo-Mora E, Cunningham M, Darboe S, Day NPJ, Luca MD, Dokova K, Dramowski A, Dunachie SJ, Bich TD, Eckmanns T, Eibach D, Emami A, Feasey N, Fisher-Pearson N, Forrest K, Garcia C, Garrett D, Gastmeier P, Giref AZ, Greer RC, Gupta V, Haller S, Haselbeck A, Hay SI, Holm M, Hopkins S, Hsia Y, Iregbu KC, Jacobs J, Jarovsky D, Javanmardi F, Jenney AWJ, Khorana M, Khusuwan S, Kissoon N, Kobeissi E, Kostyanev T, Krapp F, Krumkamp R, Kumar A, Kyu HH, Lim C, Lim K, Limmathurotsakul D, Loftus MJ, Lunn M, Ma J, Manoharan A, Marks F, May J, Mayxay M, Mturi N, Munera-Huertas T, Musicha P, Musila LA, Mussi-Pinhata MM, Naidu RN, Nakamura T, Nanavati R, Nangia S, Newton P, Ngoun C, Novotney A, Nwakanma D, Obiero CW, Ochoa TJ, Olivas-Martinez A, Olliaro P, Ooko E, Ortiz-Brizuela E, Ounchanum P, Pak GD, Paredes JL, Peleg AY, Perrone C, Phe T, Phommasone K, Plakkal N, Ponce-de-Leon A, Raad M, Ramdin T, Rattanavong S, Riddell A, Roberts T, Robotham JV, Roca A, Rosenthal VD, Rudd KE, Russell N, Sader HS, Saengchan W, Schnall J, Scott JAG, Seekaew S, Sharland M, Shivamallappa M, Sifuentes-Osornio J, Simpson AJ, Steenkeste N, Stewardson AJ, Stoeva T, Tasak N, Thaiprakong A, Thwaites G, Tigoi C, Turner C, Turner P, Doorn HR van, Velaphi S, Vongpradith A, Vongsouvath M, Vu H, Walsh T, Walson JL, Waner S, Wangrangsimakul T, Wannapinij P, Wozniak T, Sharma TEMWY, Yu KC, Zheng P, Sartorius B, Lopez AD, Stergachis A, Moore C, Dolecek C, Naghavi M. 2022. Global burden of bacterial antimicrobial resistance in 2019: a systematic analysis. The Lancet 399:629–655.

2. Kaderabkova N, Bharathwaj M, Furniss RCD, Gonzalez D, Palmer T, Mavridou DAI. 2022. The biogenesis of β-lactamase enzymes. Microbiology 168:001217.

3. Bush K, Bradford PA. 2020. Epidemiology of β-Lactamase-Producing Pathogens. Clin Microbiol Rev 33:e00047–19.

4. Fatima H, Goel N, Sinha R, Khare SK. 2021. Recent strategies for inhibiting multidrug-resistant and β-lactamase producing bacteria: A review. Colloids Surf B Biointerfaces 205:111901.

5. McEwen SA, Collignon PJ. 2018. Antimicrobial Resistance: a One Health Perspective. Microbiol Spectr 6:10.1128/microbiolspec.arba-0009–2017.

6. López-Hernández I, López-Cerero L, Fernández-Cuenca F, Pascual Á. 2022. The role of the microbiology laboratory in the diagnosis of multidrug-resistant Gram-negative bacilli infections. The importance of the determination of resistance mechanisms. Med Intensiva Engl Ed 46:455– 464.

7. Bonnin RA, Jousset AB, Emeraud C, Oueslati S, Dortet L, Naas T. 2021. Genetic Diversity, Biochemical Properties, and Detection Methods of Minor Carbapenemases in Enterobacterales. Front Med 7.

8. Guitor AK, Raphenya AR, Klunk J, Kuch M, Alcock B, Surette MG, McArthur AG, Poinar HN, Wright GD. 2019. Capturing the Resistome: a Targeted Capture Method To Reveal Antibiotic Resistance Determinants in Metagenomes. Antimicrob Agents Chemother 64:e01324–19.

9. Crofts TS, Gasparrini AJ, Dantas G. 2017. Next-generation approaches to understand and combat the antibiotic resistome. Nat Rev Microbiol 15:422–434.

10. Kantele A, Kuenzli E, Dunn SJ, Dance DAB, Newton PN, Davong V, Mero S, Pakkanen SH, Neumayr A, Hatz C, Snaith A, Kallonen T, Corander J, McNally A. 2021. Dynamics of intestinal multidrug-resistant bacteria colonisation contracted by visitors to a high-endemic setting: a prospective, daily, real-time sampling study. Lancet Microbe 2:e151–e158.

11. Osterblad M, Leistevuo T, Huovinen P. 1995. Screening for antimicrobial resistance in fecal samples by the replica plating method. J Clin Microbiol 33:3146–3149.

12. Tufic-Garutti SDS, Ramalho JVAR, Longo LG de A, de Oliveira GC, Rocha GT, Vilar LC, Dias da Costa M, Picão RC, Girão VB de C, Santoro-Lopes G, Moreira BM, Rodrigues KM de P. 2021. Acquisition of antimicrobial resistance determinants in Enterobacterales by international travelers from a large urban setting in Brazil. Travel Med Infect Dis 41:102028.

13. Fontana HYY. 2019. Occurrence and molecular characterization of AmpC-producing Extraintestinal Pathogenic Escherichia coli from human infections. Master’s thesis. UFRJ, Rio de Janeiro, Brazil.

14. Paschoal RP, Campana EH, Corrêa LL, Montezzi LF, Barrueto LRL, da Silva IR, Bonelli RR, Castro L de S, Picão RC. 2017. Concentration and Variety of Carbapenemase Producers in Recreational Coastal Waters Showing Distinct Levels of Pollution. Antimicrob Agents Chemother 61.

15. Clinical and Laboratory Standards Institute (CLSI). 2023. Performance Standards for Antimicrobial Susceptibility Testing. CLSI Supplement M100. Clinical and Laboratory Standards Institute.

16. Pérez-Pérez FJ, Hanson ND. 2002. Detection of Plasmid-Mediated AmpC β-Lactamase Genes in Clinical Isolates by Using Multiplex PCR. J Clin Microbiol 40:2153–2162.

17. Creighton J, Tibbs C. 2017. Evaluation of the MAST indirect carbapenemase test and comparison with a modified carbapenem inactivation method for the detection of carbapenemase enzymes in Gram-negative bacteria. N Z J Med Lab Sci 71:136–140.

18. Howard JC, Creighton J, Ikram R, Werno AM. 2020. Comparison of the performance of three variations of the Carbapenem Inactivation Method (CIM, modified CIM [mCIM] and in-house method (iCIM)) for the detection of carbapenemase-producing Enterobacterales and non-fermenters. J Glob Antimicrob Resist 21:78–82.

19. Tsai Y-M, Wang S, Chiu H-C, Kao C-Y, Wen L-L. 2020. Combination of modified carbapenem inactivation method (mCIM) and EDTA-CIM (eCIM) for phenotypic detection of carbapenemase-producing Enterobacteriaceae. BMC Microbiol 20:315.

20. Poirel L, Walsh TR, Cuvillier V, Nordmann P. 2011. Multiplex PCR for detection of acquired carbapenemase genes. Diagn Microbiol Infect Dis 70:119–123.

